# Identifying ancient antibiotic resistance genes in archaeological dental calculus

**DOI:** 10.1101/2024.09.23.614435

**Authors:** Francesca J. Standeven, Gwyn Dahlquist-Axe, Camilla F. Speller, Andrew Tedder, Conor J. Meehan

## Abstract

Research on ancient antimicrobial resistance is limited, and appropriate screening criteria for identifying antibiotic (ARGs) and metal resistance genes (MRGs) in archaeological samples are unclear. We assessed the impact of DNA damage and contamination on ARG and MRG detection in ancient metagenomic sequences. Starting from a set of modern oral metagenomic samples, we simulated diagenetic DNA damage as expected in ancient oral metagenomic samples. Then we estimated the impact of this damage on ARG and MRG prediction at different identity thresholds. We also examined 25 post-industrial (ca. 1850 – 1901) dental calculus samples before and after decontamination to study the rates of false positive (FP) and negative (FN) ARG and MRG predictions introduced by sample contamination. The tests showed that diagenetic damage does not significantly affect resistance gene detection, but contamination does. Furthermore, while high thresholds are advisable when feasible, overall identity thresholds do not significantly affect the rates of FPs and FNs. Additionally, comparing post-industrial and modern dental calculus revealed Tetracycline ARGs as dominant in both contaminated ancient samples and modern samples, and MLS (Macrolide, Lincosamide, and Streptogramins) ARGs as prevalent in historical samples before widespread antibiotic use.

**Data summary:** The simulated data were generated from 182 human oral biofilm samples, retrieved from the European Nucleotide Archive (ENA project: PRJNA817430) (Anderson et al., 2023). Additionally, real ancient (PRJEB1716 and PRJEB12831) and modern (PRJEB1716) metagenomic sequences were selected from metagenomic datasets published by Standeven et al. (2024).

**Impact statement:** Antimicrobial resistance (AMR) is a global health crisis. Studying the adaptability of microorganisms over centuries allows us to understand key factors that contribute to the survival and spread of antibiotic-resistant bacteria today. We know that antibiotic abuse is a key driver of AMR; however, further study into specific environmental niches that promote the evolution of antibiotic-resistant bacteria is important. For example, the extent to which the oral microbiome facilitates the increase of certain antibiotic-resistant genes and the impact of metal pollution on the spread of AMR. To investigate these key areas, it is essential to examine oral microbiomes across time, providing a complete perspective on the evolution of AMR. However, ancient metagenomics poses problems for the screening of antibiotic and metal-resistant genes in ancient bacterial DNA due to nucleotide base damage and short-read data. Through thorough threshold experimentation to establish optimal screening criteria for ancient resistance gene identification, and by addressing gaps in knowledge of ancient resistance genes, this research offers clinical significance to existing research and contributes to the development of strategies aimed at easing the impact of AMR on public health.

## Introduction

Antimicrobial resistance (AMR), declared a top global health threat by the World Health Organisation (Organisation 2020), can be described as the evolution and adaptability of microorganisms in stressful situations to improve their chances of survival. Stressful circumstances might include threats from other microorganisms including bacteria, viruses, phages, and parasites, as well as antibiotic compounds produced by plants or metals. Antibiotic resistance is often caused by an increase in antibiotic load in the environment due to antibiotic misuse, overuse, and underuse of antibiotics (Bombaywala et al. 2021) that cause bacteria to evolve, adapt, and resist drugs created to reduce their spread or destroy them. This process involves the acquisition of various antibiotic resistance genes (ARGs), or metal resistance genes (MRGs), that allow bacteria to defend themselves against certain compounds. Resistance can arise by horizontal gene transfer (HGT), which is a major cause of AMR, or from mutations that result in the acquisition of resistance-conducting genes (von Wintersdorff et al. 2016). There are different types of resistance genes, including acquired (Arzanlou et al. 2017), intrinsic (Blair et al. 2015), proto (Singh et al. 2019), and silent resistance (Singh et al. 2019).

The use of databases and consistent classification of ARGs is essential (Alcock *et al*. 2019), and various bioinformatics software, databases, and data-sharing resources are available for identifying, predicting, cataloguing, and analysing molecular sequences of ARGs (McArthur and Tsang 2017). Various tools to detect ARGs from metagenomic sequences include AMR++ (Bonin et al. 2023), CARD RGI (Alcock et al. 2023), ARG-ANNOT (Gupta et al. 2014), ResFinder (Bortolaia et al. 2020), and Meta-MARC (Lakin et al. 2019). There are also tools made just for identifying metal and biocide-resistant MRGs such as BacMet (Pal et al. 2014); tools such as the AMR++ pipeline can screen for both ARGs and MRGs. These pipelines all typically identify resistance genes by aligning DNA sequences or whole-genomes to reference databases (Zankari et al. 2017; Alcock et al. 2019; Feldgarden et al. 2019; Bortolaia et al. 2020; Feldgarden et al. 2021; Florensa et al. 2022; NCBI 2024) to detect genes and single nucleotide polymorphisms (SNPs). Tools that access a combination of these databases, like AMR++, can help reduce the impact of database bias.

Microbial DNA recovered from archaeological or historical samples has a high potential to inform ARG and MRG studies by providing a perspective into resistance gene prevalence before widespread antibiotic use. However, there is little guidance within bioarcheology on the right tools and databases to use for ancient ARG and MRG analyses. However, some tools are known to perform well on DNA with ancient traits and some researchers may prefer pipelines like AMR++ and ARG-ANNOT, which work effectively with short sequences. To elaborate, ancient microbial DNA analysis is fraught with challenges, including poor quality, limited quantity, and high contamination levels of ancient DNA (Gaeta 2021). aDNA is typically damaged through common forms of chemical processes that degrade DNA over long periods such as depurination, which causes changes in nucleotide base pairs (Dabney et al. 2013) and can also cause fragmentation (Kistler et al. 2017). These structural issues of aDNA have the potential to hinder bioinformatic tools from aligning nucleotides to reference genomes, but it is not known exactly to what extent. Regarding contamination, it is strongly advised that environmental and laboratory blanks decontaminate samples before screening for resistance genes following research on ARGs identified in mummified remains (Santiago-Rodriguez et al. 2018) that sparked criticism of the lack of decontamination protocols (Eisenhofer and Weyrich 2018). Environmental decontamination is crucial because archaeological skeletons can remain *in situ* for thousands of years, accumulating contaminant DNA from the environment that may accrue similar damage patterns to the endogenous DNA. Due to the widespread distribution of ARGs across landscapes, the surrounding soil contains genetic material that was not originally part of the ancient remains.

Bioinformaticians also encounter challenges when determining suitable cut-off thresholds such as e-values and sequence identity (Ji et al. 2022), and appropriate threshold selection for ancient ARG identification is especially difficult. This is demonstrated in a study by Rascovan et al. (2016) that explores ARGs in several previously published ancient metagenomic datasets. The authors use both stringent (>90% similarity) and non-stringent (*evalue* <1E^-10^) parameters to screen for ARGs against the ARGANNOT database and highlighted that three more groups of ARGs were identified using less strict thresholds. The authors are aware that using less stringent parameters increases the risk of false positives (FPs), but they argue that stricter thresholds pose an even greater risk of false negatives (FNs). Except for the study by Rascovan et al. (2016), there is a lack of discussion regarding the appropriate thresholds for analysing resistance genes in ancient samples.

Ancient resistance genes have been found in arctic soil (Perron et al. 2015), caves (Bhullar et al. 2012; Pawlowski et al. 2016), and permafrost (D’Costa et al. 2011; Kashuba et al. 2017). Although the study of ARGs is intriguing in these contexts, the concept of their antiquity is not surprising (Relman and Lipsitch 2018; Kraemer et al. 2019) because bacteria have always developed defence mechanisms to survive (Baron et al. 2018). Antibiotics are naturally occurring (Van Goethem et al. 2018) and most antibiotics today are derived from natural sources (Hutchings et al. 2019).

Ancient resistance genes in human microbiomes are intriguing because delving into their ancient presence before the influence of extreme selective pressure allows us to track evolutionary traits and uncover novel mechanisms of bacteria in clean models. For example, Rascovan et al. (2016) examined 10^th^ and 12^th^-century dental calculus and found a new class B Beta-lactamase family showing high penicillinase activity. Ancient dental calculus (fossilized dental plaque) is a promising source of resistance genes because they have been shown to entrap oral microorganisms and their genes and preserve them over thousands of years (Warinner et al. 2014; Weyrich et al. 2017; Hendy et al. 2018; Fotakis et al. 2020; Neukamm et al. 2020).

Examining resistance genes in archaeological contexts helps us understand how significant lifestyle changes, such as industrialisation, may have influenced the spread and prevalence of specific resistance genes in human oral microbiomes. For example, smoke pollution in Manchester, UK, during the Industrial Revolution was particularly severe, and the poorest inhabitants were compelled to dwell amid the mills and factories in the north-easterly regions, while some better-paid workers may have avoided the worst of the pollution (Mosley 2008). Such a significant historical transition as this would likely increase the levels of MRGs in the environment. It is also noteworthy that heavy metals are shown to operate as co-selecting agents in the spread of antibiotic resistance in human pathogens across various environments (Nguyen et al. 2019). Moreover, the coexistence of ARGs, metal resistance genes (MRGs), and mobile genetic elements in resistomes of modern dental plaque has indicated the existence of a co-selection process in the oral microbiome (Kang et al. 2021). Given this evidence, it is worthwhile to explore the role of heavy metals in the evolution of ARGs in Industrial Age oral microbiomes that were heavily exposed to metal pollution and untouched by modern prescription antibiotics.

Research on ARGs in ancient human microbiomes is still scarce due to the limited number of published ancient ARGs identified in dental calculus. Additionally, there are no established guidelines for their discovery, as the impact of ancient DNA characteristics on ARG identification has not been comprehensively studied. Therefore, further research is needed to address these questions and fill other gaps in our understanding of resistance genes, such as the natural prevalence of resistance genes in oral microbiomes. This study explores how environmental contamination and damage resulting from diagenesis affects the discovery and analysis of resistance genes in ancient samples. Moreover, we analyze dental calculus samples from different historic periods to investigate which resistance genes in the oral microbiome can be correlated with historical human transitions such as industrialisation and modern medical innovations.

## Materials and Methods

### Quantifying the effects of diagenetic damage

To understand the impact of diagenetic damage on ARG identification, we first carried out a threshold experiment using simulated ancient DNA. First, we downloaded 182 public human biofilm samples through the European Nucleotide Archive (ENA) from modern oral microbiomes that were healthy, caries-active, or periodontally diseased, collected from the University Hospital in Heidelberg, Germany (ENA project title: PRJNA817430) (Anderson et al. 2023). To simulate ‘damaged’ versions of these 182 samples, the fastq files were converted into fasta format and then the ‘deamSim’ function from gargammel v1.1.4 (Renaud et al. 2017) was applied with the following default parameters-damage 0.03, 0.4, 0.01, 0.7 to deaminate nucleotide bases. Deaminated reads were then recombined with their initial per base quality scores to recreate fastq paired-end read files.

Having a batch of modern DNA and a batch of ‘damaged’ DNA to simulate aDNA enabled us to calculate true positive (TP), false positive (FP), true negative (TN), and false negative (FN) in our ARGs data. To do this, we implemented Nextflow v23.04.1 to run the MEGARes and AMR++ pipeline v3 (Bonin et al. 2023) first on the modern samples using three sequence identity percentage parameters: low (25%), medium (50%), and high (75%), and then on the simulated aDNA with the same thresholds. We used the modern results at a 75% parameter as the ground truth.

Following AMR++, an analysis of variance (ANOVA) test was performed in R to measure significant differences of resistance gene counts between ancient and modern threshold groups. Then, the type of individual resistance genes between groups (based on unique MEGARes accession numbers) was analysed by counting the number of times each MEGARes accession code (ARGs or MRGs) was called in each group. This data was used to count the number of TPs, FPs, TNs, and FNs based on comparison to the modern 75% sequence identity threshold results. To take a closer look at the small differences between the types of ARGs being produced between these groups, resistance gene group types (as opposed to just the single gene identified through a MEGARes accession) were assembled, counted across the threshold groups, and an ANOVA test was performed to show statistical differences in gene type across groups.

### Contamination experiment

Next, we sought to determine the degree to which microbial contamination from the environment influenced the recovery of ARGs and MRGs in historical samples, specifically the level of false positives introduced due to modern contamination of samples. For this experiment, we used metagenomic data from 25 post-industrial (ca. 1850 – 1901) dental calculus samples (50 paired-end fastq files) from varying sites across the UK (Table **S1**). Also included in this study are 10 modern (21^st^ Century) oral metagenomic samples from the University of Tennessee, USA, (20 paired-end fastq files) which are involved in a separate experiment outlined below. Detailed information on these metagenomic datasets is reported in Standeven et al. 2024.

All metagenomic datasets went through a series of pre-processing steps including quality control, decontamination, and damage authentication, which can be found in more detail in the data publication by Standeven et al. 2024. To quality check samples, raw fastq read data were analysed using FastQC v0.11.9 with default parameters (Babraham Bioinformatics 2019). Then, Fastp v0.23.2 (Chen *et al*. 2018) was used with default parameters to filter and trim poor-quality reads, cut adapters, repair mismatched base pairs, and ensure overall quality. BBduk (Joint Genome Institute 2023) was utilised with default parameters to decontaminate samples by filtering out homologous sequences found in paired environmental and laboratory blanks. To authenticate ancient sequences, we retained human reads from outputs produced by Centrifuge v1.0.3 (Kim et al. 2016) and used seqtk ‘subseq’ to convert them into fastq files which were then mapped to the human genome (hg38) (NCBI 2013) using BWA mem v0.7.17 (Li and Durbin 2009). SAMtools v1.12 (Danecek et al. 2021) (-view -rmdup -flagstat -sort -index) was used to format the alignments and sort into BAM files which were run through mapDamage2 v2.2.2 with default parameters (Jónsson et al. 2013).

The AMR++ pipeline was applied to these 25 post-industrial samples before and after decontamination, producing two separate resistance gene results. Firstly, we compared the number of resistance genes before and after decontamination; a list of counts per resistance gene in each group was also created for comparison. Then, as described above, an ANOVA test was performed in R to show statistical differences in types of resistance genes between the contaminated and decontaminated group to calculate TPs, FPs, TNs, and FNs, using the decontaminated results as the ground truth. We also identified which resistant genes are responsible for significant differences between the two groups, and R was used to list which MEGARes numbers were unique to both groups.

### Post-industrial versus modern analysis

The discovery and mass manufacture of antibiotics changed the way humans experience AMR today – it is no longer solely a microbial conflict taking place discreetly in the environment, but also a large-scale issue affecting human health and lifestyle. Our study also investigated which genes in particular have driven that transformation and continue to impact the oral microbiome. Ten modern dental calculus samples (**Table S1**) were put through the AMR++ pipeline and resistance genes were compared with the post-industrial results. Since we chose a 50% sequence identity threshold for our ancient data, this parameter was also applied to our modern data to make the results comparable.

Percentages of ARGs and MRGs were calculated in each group and compared to highlight resistance genes associated with ancient and modern oral microbiomes, or any increases or declines in genes across the two periods. To calculate ARGs and MRGs coverage in these groups, the MEGARes drugs database was downloaded, and an in-house Python script was made to extract reference genes in fasta format. BWA mem v0.7.17 (Li and Durbin 2009) (-v 3 -t 20) was then utilised to align resistance query genes to reference genes, and then SAMtools v1.12 (Danecek et al. 2021) (sort --threads 20 coverage) was used to generate coverage tables.

## Results

### The impact of damage on ARG prediction

ANOVA was conducted to compare the effect of DNA damage on predicted counts of resistance genes (MEGARes accession numbers). No significant differences were found between the simulated ancient and modern threshold groups, F value = 0.917, p = 0.469. A table of counts per resistance gene for each ancient and modern threshold group was produced (**Table S2**), showing that slightly more than half of individual genes (51%) do not exhibit differences in counts across the specified modern and ancient threshold groups. From these data, the TP, FP, TN, and FN percentages were calculated (**Figure 1; Table S3**). The combination of TP and TN percentages in each group is close to the ground truth group (modern 75%). FN percentages are low (<15%) and restricted to ancient groups, and the FP proportion is higher than the ground truth (10%–45%) in all groups.

**Figure 1.**
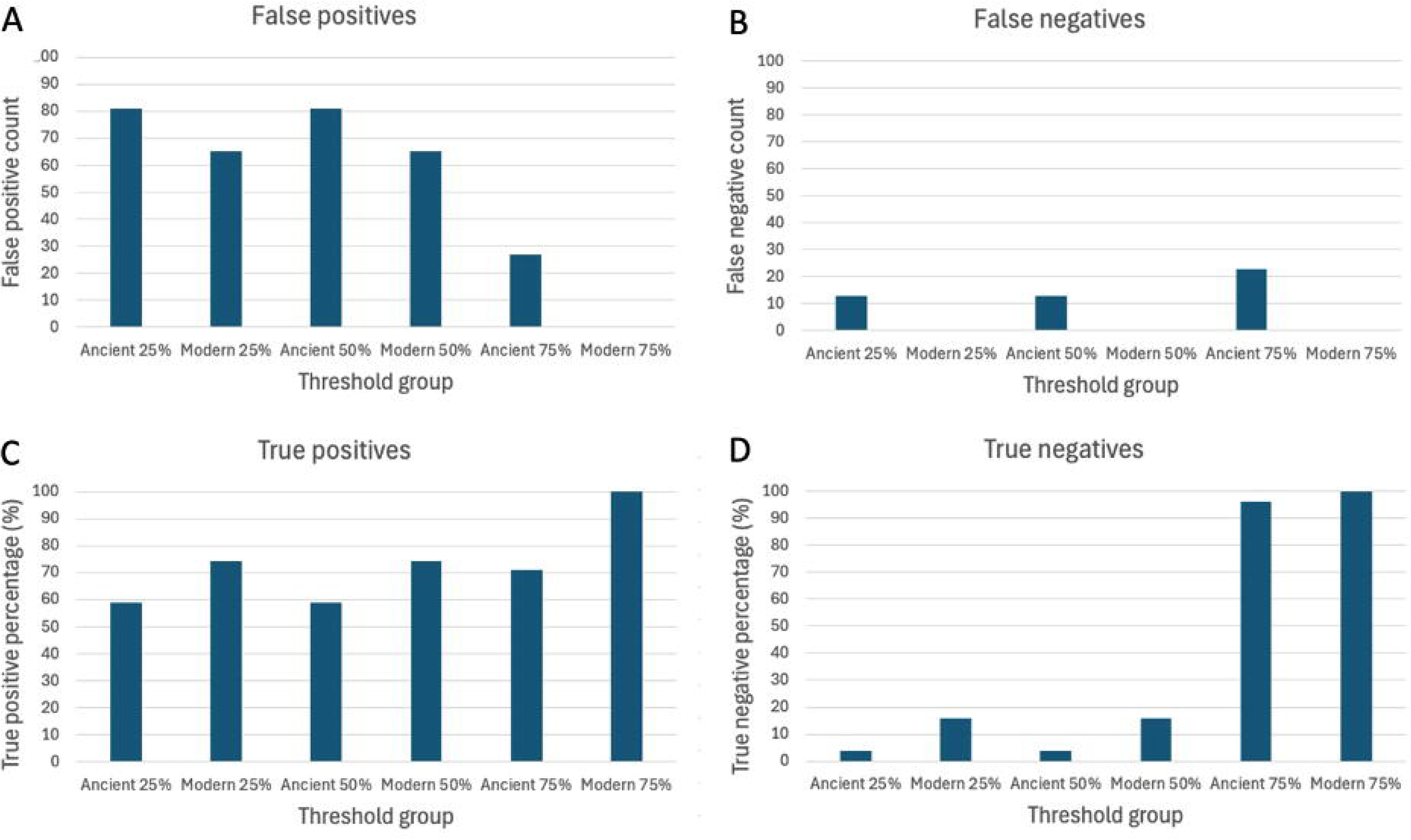
(a) False positive (FP), (b) false negative (FN), (c) true positive (TP), and (d) true negative (TN) count and percentage per modern and ancient threshold group.

Although there are no statistically significant differences in the individual resistance prediction counts, differences in TPs, FPs, TNs, and FNs suggest that there may be differences per resistance group type. Therefore, the resistance gene group types were counted per threshold group and are displayed in a heatmap (**Figure 2; Table S4**). ANOVA was performed to see statistical differences between resistance gene group type counts across the different threshold groups and no significant differences were found, F value = 0.337, p = 0.89. Many group type counts are consistent across thresholds like Tetracycline, Sulfonamides, Spiropyrimidinetriones, etc. However, upon closer inspection, these findings show that certain drug-resistant categories are not reliable under particular thresholds as they are likely to call more FPs. For example, when looking for Copper, Iron, or Pleuromutilin resistance, **Figure 2** shows that these categories of drug resistance do not exist in the 75% ancient and modern threshold groups, the latter being our basis for the ground truth in this analysis.

**Figure 2.**
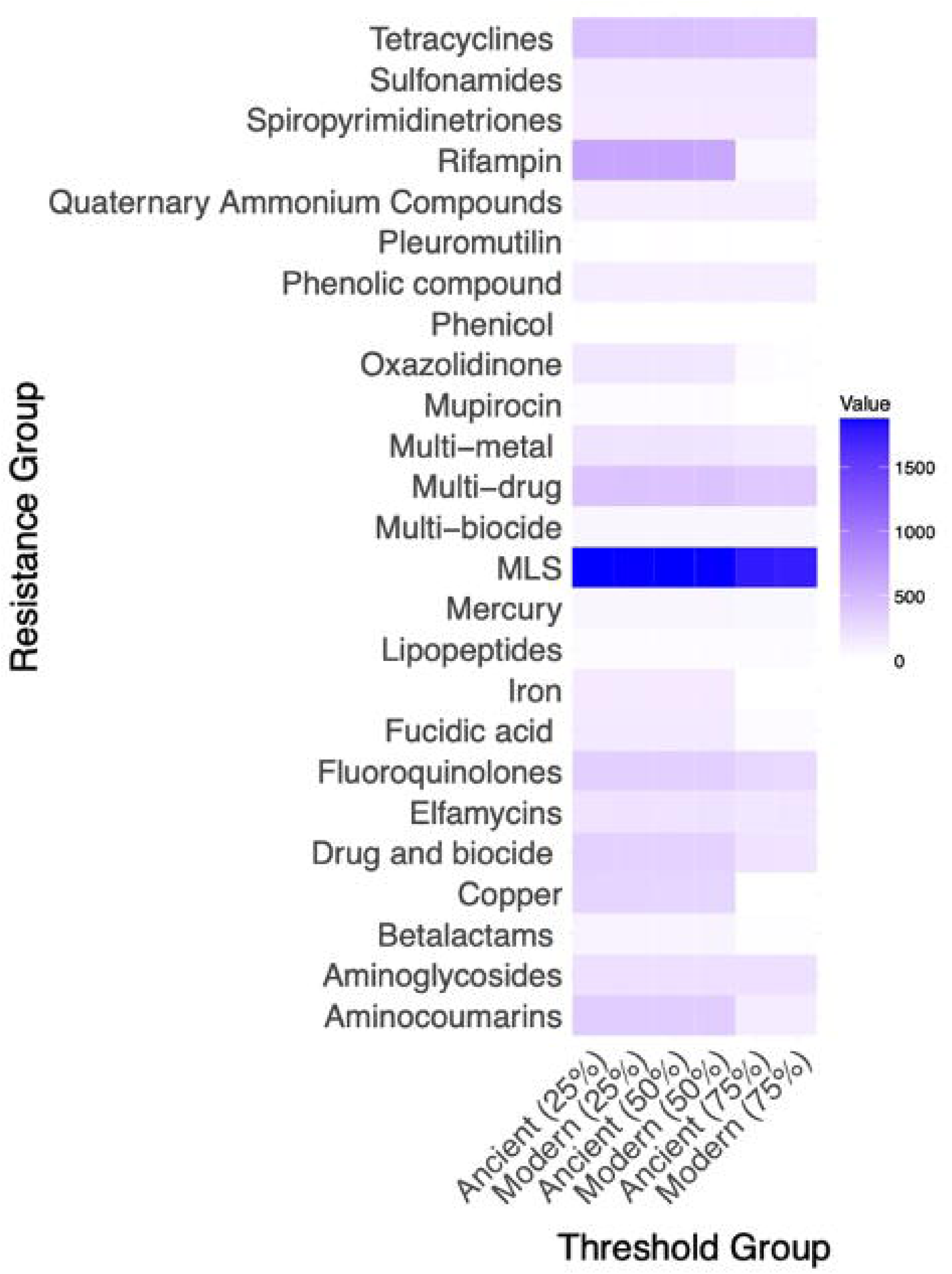
The colour gradient key on the heatmap represents the count of reads that are mapped to a specific gene in a given threshold group, ranging from white for 0 counts to light purple for counts up to 500. As the count increases, the colour transitions to dark purple at 1000 counts and shifts to dark blue beyond 1500 counts. The gradient shows the increasing abundance of genes in across groups, with lighter colours corresponding to lower counts and darker reflecting higher counts.

### The impact of contamination

For real ancient data, the ‘runsnp’ process in the pipeline failed for thresholds of 75% and 50%, likely due to data quality issues in the fastq files, such as low-quality reads. The pipeline ran without SNP errors at a 25% threshold. Since the main focus of this analysis is on resistance gene categories rather than an in-depth SNP analysis, the SNP step is not critical. Achieving results as close to the ground truth as possible by using a higher threshold is preferred. However, the pipeline at 75% spent more CPU time on failed tasks compared to 50%, thus, a 50% threshold was selected as a compromise.

Resistance genes were counted in contaminated and decontaminated groups and compared (**Table S5**) to determine significant variation in the resistance gene count between the groups. A total of 1,286 individual genes were counted in the contaminated group and 111 were counted in the decontamination group, a significant difference in gene counts between the groups (p-value < 2e-16). The resistance gene counts were then used to calculate TPs, FPs, TNs, and FNs, using the decontaminated results as the ground truth (**Table S6**). The decontaminated group contained all the TPs, however, the exact counts for each gene were only correct in three instances (MEG_3838, MEG_5405, MEG_5776). The contaminated group contained 198 FPs.

The decontaminated resistance gene list was removed from the original contaminated list for each sample and what remained was 177 unique MEGARes accession numbers (different to counts) unique to the contaminated group (**Table S7**). The percentage of different resistance gene types identified in the contaminated and decontaminated groups is shown in **Fig 3** (**Table S8**). Although certain MEGARes accession numbers are unique to the contaminated group, **Fig 3** shows that some resistance gene categories that these genes belong to are not. For example, the contaminated group has many unique Betalactam and Aminoglycosides ARGs that do not exist in the decontaminated group, but this is not to say that all Betalactam and Aminoglycosides ARGs are contaminants because the decontaminated group also have a few Betalactam and Aminoglycosides ARGs.

**Figure 3.**
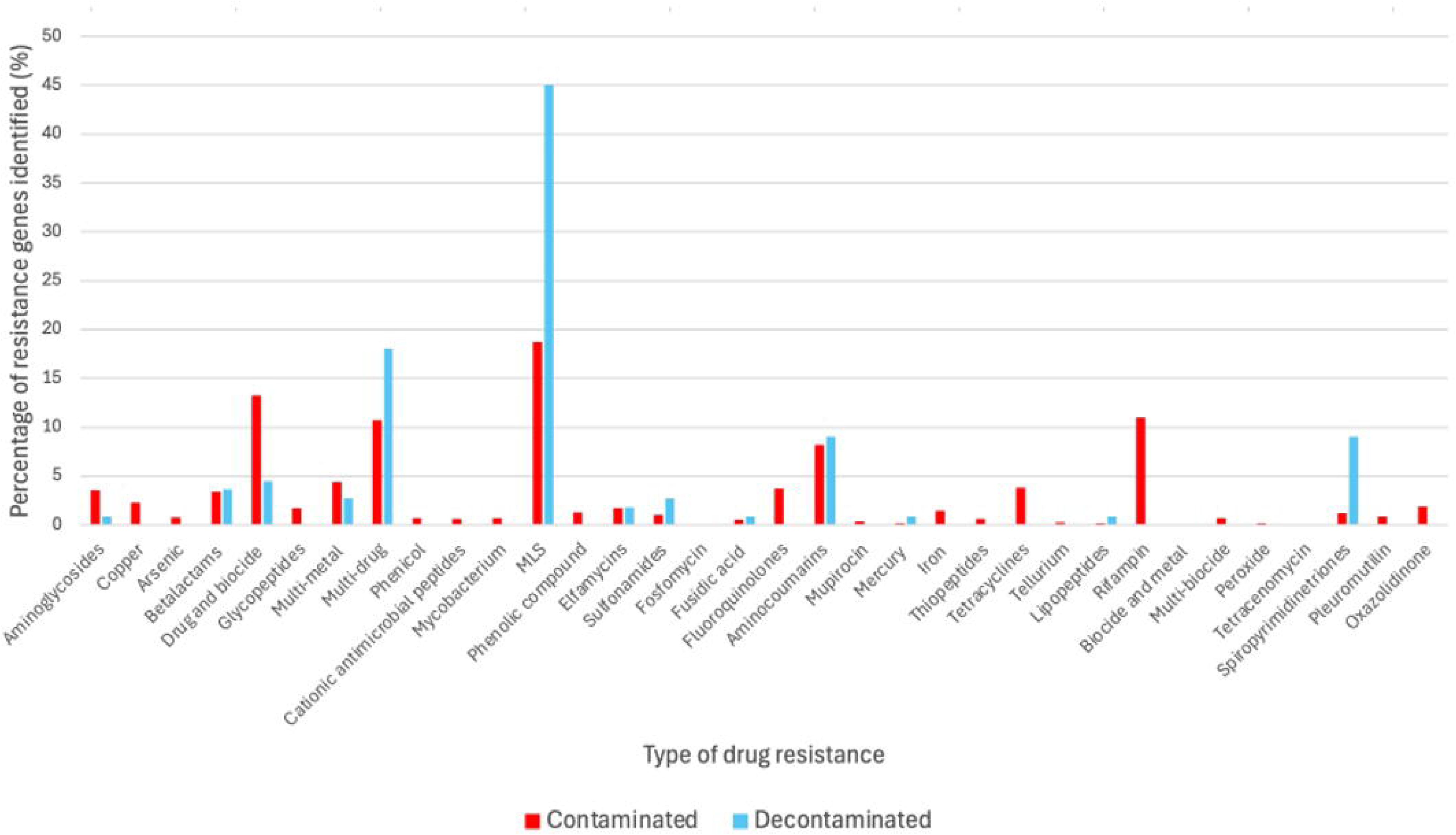
Percentage of putative resistance genes identified in contaminated and decontaminated subgroups.

On the other hand, there are entire drug resistance categories in the contaminated group that are completely absent in the decontamination samples and are likely to be an artefact of modern contamination: Aminocoumarins; Arsenic; Biocides; Cationic antimicrobial peptides; Copper; Fluoroquinolones; Fosfomycin; Glycopeptide; Iron; Phenicol; *Mycobacterium tuberculosis*-specific drugs; Oxazolidinone; Pleuromutilin; Rifampin; Tellurium; Thiopeptides; Tetracyclines; and Tetracenomycin. These findings show that contamination has a large impact on the retrieval of ARGs and MRGs in ancient samples that have not been decontaminated.

### Patterns in modern and ancient resistance genes

The percentage of different resistance gene types identified in ancient (post-industrial) and modern dental calculus samples is shown in **Figure 4**. The ancient dataset has 15 more samples than the modern dataset, so direct similarity comparisons cannot be made. However, it is clear that certain resistance gene types are specific to certain periods. The ancient oral microbiome revealed the following resistance genes: including MLS (Macrolide, Lincosamide, and Streptogramins), Beta-lactams, Fusidic acid, Multi-metal, Elfamycins, Mercury, Multi-drug, Sulfonamides, Spiropyrimidinetriones, and Aminocoumarins. Several resistance genes are restricted to modern samples: Aminoglycosides, Copper, Fluoroquinolones, Iron, Lipopeptides, Oxazolidinone, Rifampin, and Tetracyclines. Resistance to MLS (43.9%) are ARGs heavily linked to the ancient oral microbiome and are less common in modern oral microbiota (22.55%). Instead, resistance to Beta-lactams (10.78%) and Tetracyclines (48.04%) are the dominant resistance genes in the modern oral microbiomes within this study. **Figure 5** and **Table S9** show overall good coverage (%) for ARGs and MRGs in modern and post-industrial groups.

**Figure 4.**
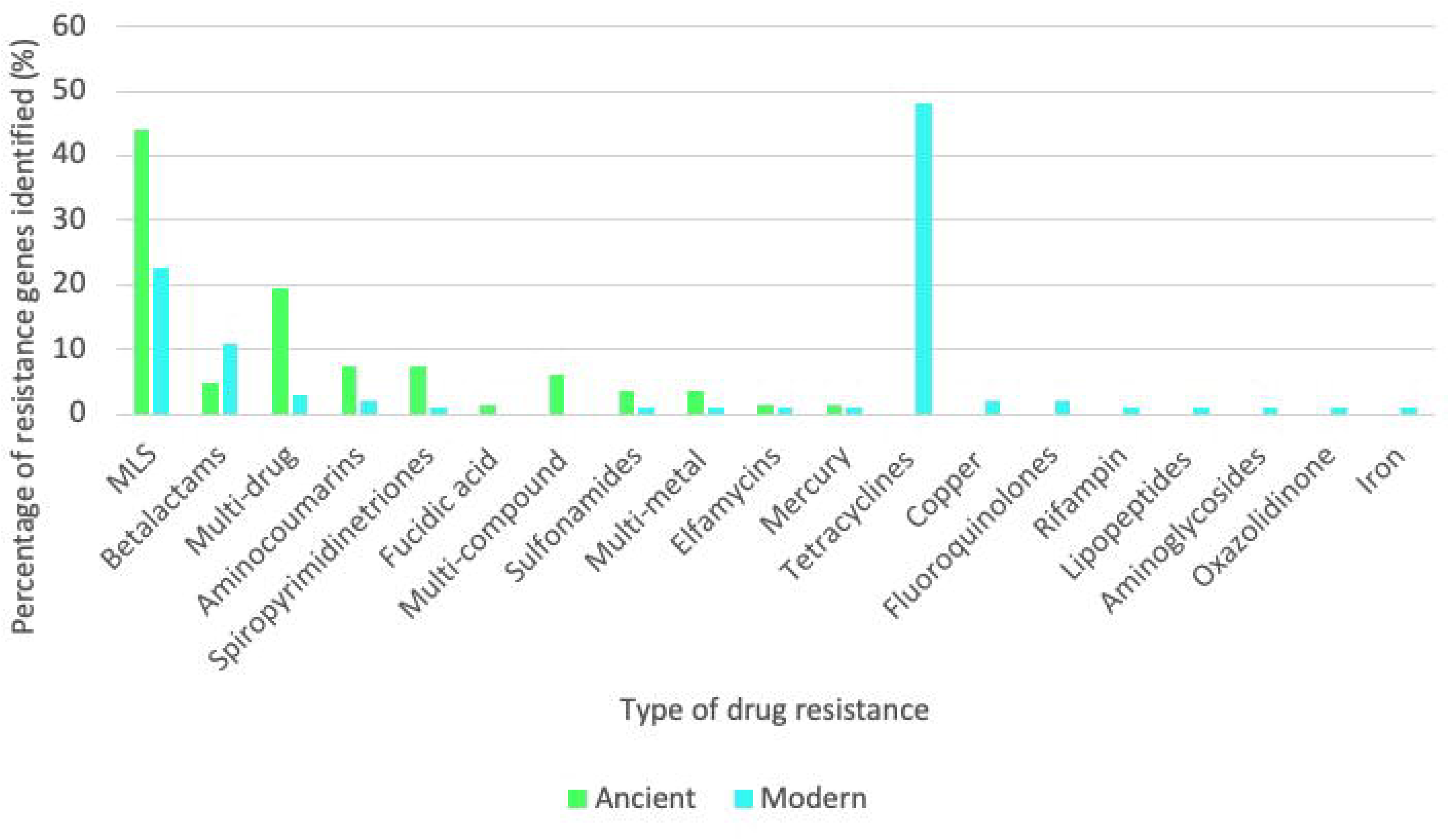
Percentage of putative resistance genes identified in post-industrial and modern oral resistomes.

**Figure 5.**
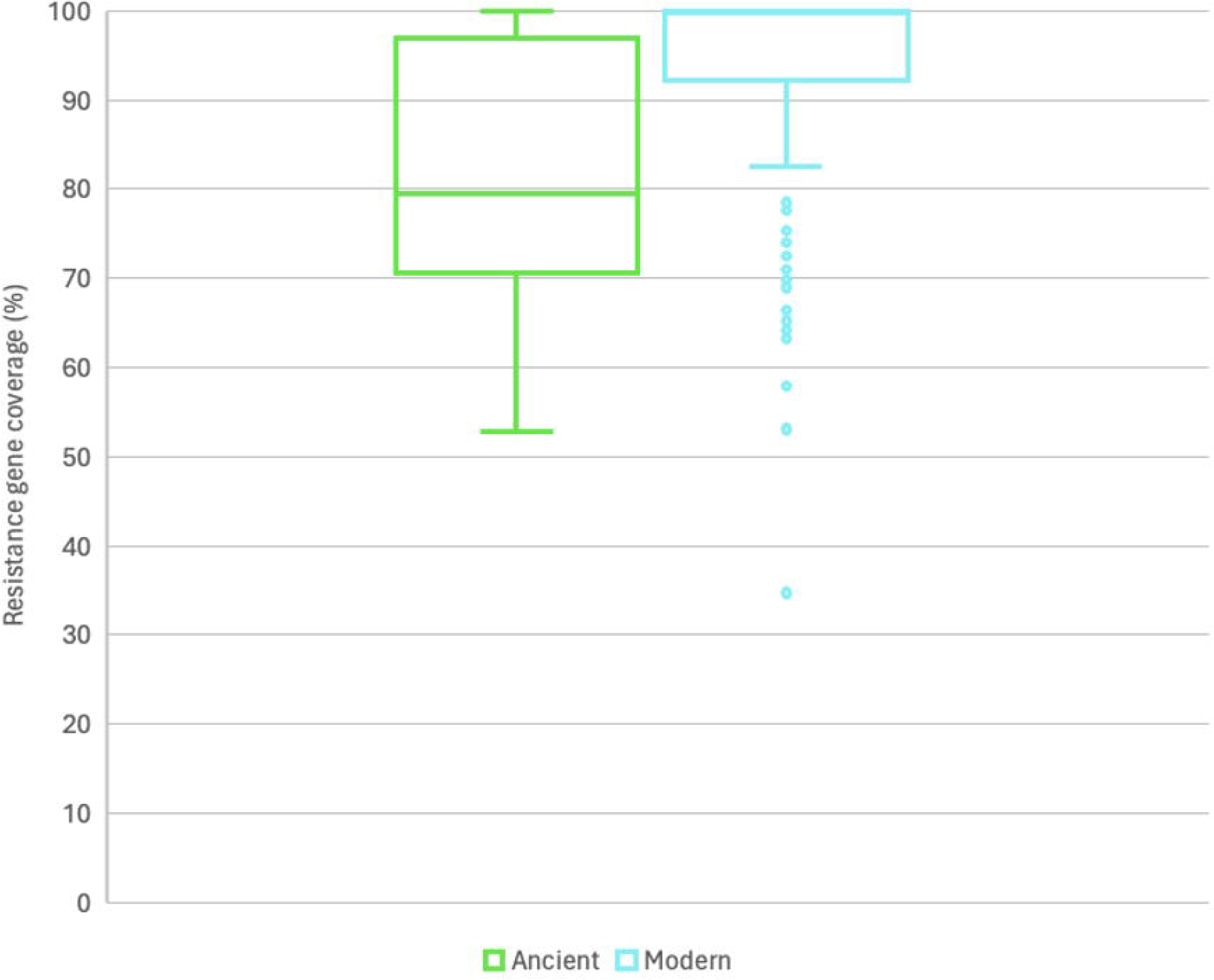
Coverage (%) for resistance genes in ancient and modern groups.

## Discussion

This research investigated factors affecting the retrieval of resistance genes from ancient samples, focusing on damage and contamination. Through simulated data, it was found that gene counts and types remained statistically consistent between ancient and modern threshold groups, indicating minimal impact from damage (deamination). However, false positives and negatives in ancient samples necessitate caution. Contamination, on the other hand, significantly affected gene retrieval, with a notable decrease in gene counts and type post-decontamination. Analysis of post-industrial and modern dental calculus revealed a prominent peak of Tetracycline resistance genes in modern oral microbiomes, absent in ancient samples, indicating evidence of widespread use of Tetracycline following the post-industrial period into the modern era and its clear impact on the oral microbiome.

### Minimal effect of damage on resistance gene retrieval

It could be predicted that lowering the sequence identity barrier would increase resistance gene counts between groups since loose cut-offs allow for the detection of more DNA sequences. The ANOVA test found no significant difference in resistance gene count between simulated ancient and modern threshold groups. This finding is informative since it indicates that the percentage of FPs is not very high when lowering the identity thresholds. Nonetheless, this is based solely on overall gene count, and it does not into account the differences in resistance gene types between the groups which could still lead to misleading results. Instead of counting all of the genes simultaneously, we compared how many times each gene was counted in each group and compared them. There were no significant differences in gene type since most gene counts were consistent across each group. This is an important outcome because it shows that the pipeline detects similar types of genes in ancient and modern samples, regardless of damage or threshold.

No significant overall differences in gene count and variability between the ancient and modern threshold groups may come as a surprise as damaged DNA may have the potential to read differently to modern DNA and produce varying results. The reason for this might be because, unlike attempting to align longer and more fragmented core bacterial genomes, the smaller size of ancient variable genes with fewer nucleotides, such as resistance mechanisms with fewer nucleotides, may be less challenging to identify. Additionally, as a result of the AMR crisis, it can be argued that there is a strong research focus on ARGs and MRGs, with extensive sequencing efforts contributing to their detailed characterisation. This is a promising outcome, as it indicates that researchers can generally confidently conduct resistance gene analysis without concern that damage will distort the retrieval of resistance genes from metagenomic sequences. Nonetheless, it is critical to interpret these data with caution. We added artificial damage to modern bacterial DNA using defined parameters, but in fact, aDNA can experience several forms of degradation (Uysal Yoca et al. 2021), all of which could impact results differently. For example, aDNA that has been *in situ* for 5000 years and has undergone considerable deamination (the transformation of cytosine to uracil and interpreted as thymine) may have a very different structure to DNA that has been buried for a century and exposed to harsh acidic conditions. This merits further investigation.

Despite that statistical testing showed threshold and damage do cause statistically measurably impacts, a closer analysis of the TP, FP, TN, and FN count reveals some discrepancies between groups, indicating that some refinement of the method is needed. There were no FNs in modern groups at any threshold, whereas there were some FNs (<10%) in the ancient groups, the highest being in the ancient 75% group. This is expected because tightening the threshold allows for fewer genes to be identified, which means that some genes may be overlooked. Given that these genes are also ‘ancient’, it is expected that sequences with some damage in key portions of ARG sequences may drop the similarity to modern reference genes below the 75% threshold. In comparison to the FN proportion, the FP rate is high across all groups (between 10% and 45%), increases at lower thresholds, and is the highest among the ancient groups. In general, the relaxing of criteria allows for the identification of more genes, and unavoidably, some of those genes will not be genuine. Moreover, the high FP rate among ancient samples may be linked to the program’s ability to process and align modified nucleotides, causing false results, especially at lower thresholds (50% and 25%).

In ancient samples, low thresholds cause more FPs, and high thresholds cause more FNs. However, the FN proportion is always lower than the FP proportion in ancient samples across all thresholds. There is a large element of debate in deciding which thresholds are more appropriate to use regarding the proportion of FPs and FNs and it is down to context to conclude. Some studies have recommended using stricter sequence identity cutoffs between 80%-95% for detecting modern resistomes (Bengtsson-Palme et al. 2017; Liao et al. 2022). Other literature argues that using strict cut-offs is reliable in avoiding false positives, but it can lead to a high false negative rate. Since many ARGs do not match closely with known databases, strict similarity thresholds can cause many ARGs to be overlooked (Arango-Argoty et al. 2018). This is an important consideration in the context of aDNA. While stringent criteria are necessary for accurately identifying very short-read DNA to minimise false positives, it is essential to recognize that a more rigorous approach may inadvertently overlook lesser-known or extinct ARGs from ancient metagenomic sequences. For example, previous research on ancient ARGs has argued that while lenient cut-offs result in FPs to a certain extent, the frequency of FNs produced by tighter thresholds is more detrimental (Rascovan et al. 2016). Although there are medical situations where FNs have grave consequences, our results suggest that in archaeological contexts, the larger proportion of present untrustworthy resistance genes is more misleading than the fewer resistance genes that were missed.

The use of low thresholds on aDNA is sometimes unavoidable. This study found that SNP analysis was unsuccessful at higher thresholds (50% and 75%) in cases of low read quality in our archaeological aDNA dataset. Since our focus is on resistance categories rather than specific genetic variations, poor SNP calling did not impact the overall results of the study. However, researchers aiming to understand the development, evolution and spread of ancient resistance genes using gene variation may be affected by these threshold settings and might need to use lower thresholds, which increases the risk of false positives. Studies focusing on more effective aDNA extraction methods as well as improved sequencing coverage would help overcome this challenge.

Future research should also consider investigating additional thresholds beyond sequence identity, such as target coverage cut-offs, which focus on the proportion of the reference gene that is aligned. For instance, a study by Cooper et al. (2024) demonstrated that antimicrobial resistance genes (ARGs) can still be detected even when the target organism is present in lower amounts and the coverage ranges between 40-60%. However, detecting ARGs at these lower coverage levels may increase the risk of identifying similar but incorrect ARGs (Cooper et al. 2024). Given this finding, it is important to evaluate how these thresholds influence the detection of resistance genes in ancient metagenomic sequences.

We investigated if any specific drug categories were driving these minor discrepancies. It was revealed that Copper, Iron, and Pleuromutilin resistance are unreliable genes in our ancient groups because they are absent in the 75% ancient and modern threshold groups. Moreover, some MEG accessions of the same resistance type have different TP and FP rates. For example, one Aminocoumarin ARG is trustworthy (MEG_5380), while the rest are not (MEG_3247, MEG_3248, MEG_3249, MEG_5364, MEG_5380, MEG_5389, MEG_5396, MEG_3247, MEG_3248, and MEG_3249). These small differences are important because although the statistics suggest that using certain thresholds on ancient data are appropriate, stricter parameters are recommended when screening for certain drug-resistant categories.

### Significant effect of contamination on resistance gene retrieval

We observed that the quantity and types of resistance genes in dental calculus before and after decontamination are significantly different. This result was attributed to the vast number and type of genes unique to the contaminated group that did not exist in the decontaminated group. Seven different Tetracycline ARGs were removed from the dental calculus batch after filtering due to being false positives. This demonstrates the level of modern contamination that, if unfiltered, may be found in ancient samples and result in biased conclusions. Although not surprising given how vulnerable aDNA is to environmental contamination, this is a major finding because previous publications have expressed significant concerns around research that does not properly decontaminate aDNA (Eisenhofer and Weyrich 2018). Studies that do not use environmental (bone, tooth root, dirt, etc.) and laboratory (extraction, library) blanks to detect and remove putative contaminants may explain why they detect modern ARGs such as those conferring to Vancomycin and Tetracycline in dental calculus (Santiago-Rodriguez et al. 2018; Ottoni et al. 2021). Furthermore, Warinner *et al*. 2014 have shown the absence of Tetracycline ARGs in bacteria from ancient dental calculus, which makes sense considering the lack of selective pressure from this antibiotic in the past.

On the other hand, it is not completely implausible to suppose that ancient Tetracycline ARGs could be identified in historical remains. Ancient populations are believed to have created fermentation mixtures containing bacteria that produce Tetracycline antibiotics (Nelson et al. 2010), and where there are Tetracyclines, there are Tetracycline ARGs. Furthermore, the presence of certain drug resistance categories in contaminated samples but not in decontaminated samples does not necessarily imply their absence in ancient resistomes. This study identified a limited number of ARGs from a restricted sample set. Therefore, additional research involving a larger number of samples is required to conclusively determine whether entire categories of drug resistance, such as tetracycline resistance, are truly contaminants.

Instead of filtering out possible ARG contaminants, studies (Ottoni et al. 2021) have used systems like SourceTracker (Knights et al. 2011) to assess the risk of contamination by determining the proportion and origin of microorganisms in a sample; however, to properly reduce the impact of contamination, it should not be used alone and without decontaminating using both an environment and a laboratory blank. Overall, we argue that based on our findings, contamination (unlike damage) has a large effect on the type and amount of resistance genes being retrieved; thus, adequate decontamination protocols are required.

### The rise and emergence of specific resistance genes in oral microbiomes

Following decontamination, we were able to reliably analyse resistance genes in our post-industrial and modern oral microbiomes. Common ARGs found in people’s mouths today are Tetracycline resistance genes (Seville et al. 2009; Rôças and Siqueira 2013; Arredondo et al. 2021; Brooks et al. 2022; Vázquez-Ramos et al. 2022; Morales-Dorantes et al. 2023), and we can expect to see less of these ARGs in ancient samples before the 1930s because there was less selective pressure from antibiotics before their discovery and human manufacture. Our post-industrial versus modern dental calculus analysis has added evidence to this notion, as 48.04% of ARGs in modern samples conferred Tetracycline while none were found in ancient samples. There was also a rise in Beta-lactam ARG proportions from 4.88% in ancient samples to 10.78% in modern ones. This shows that certain genes, such as Tetracycline ARGs, are modern occurrences that have increased alarmingly inside the oral microbiome. Therefore, this information could be crucial for determining how these resistance genes spread, whether the oral microbiome is a site of transmission, or if they are spreading in other areas and the oral microbiome, like other biofilms, (Guo et al. 2018) serves as a source for their dissemination.

Worthy of discussion in the post-industrial vs modern analysis is how the presence of certain ARGs and MRGs came to be in ancient samples. Many antibiotics are ancient and produced naturally by bacteria, fungi, or metals (Tahlan and Jensen 2013; Wolański et al. 2016; Li and Seiple 2017; McDonald et al. 2017; Yarlagadda et al. 2020; Long et al. 2021; Mori and Abe 2024; Turner 2024), explaining their presence before the mass production of antibiotics following the 1920s. Other antibiotics, such as Sulfonamides and Spiropyrimidinetriones ARGs, are synthetic compounds and their existence in the environment today is a result of modern human pharmaceutical use, making their presence in our post-industrial dental calculus confusing.

To explore why certain ARGs are found in post-industrial samples, we can re-examine their occurrence in the previously mentioned contaminated dental calculus. Sulfonamide ARGs were counted 13 times in the contaminated group, and three times in the decontaminated group, and Spiropyrimidinetriones ARGs were counted 16 times in the contaminated group and 10 times in the decontaminated group. Several of these ARGs are FPs, so they are likely contaminants, but we can offer explanations for the TPs and the reasoning for their presence in decontaminated post-industrial samples. Some natural antibiotics may have similar structures to Sulfonamides or Spiropyrimidinetriones, and ancient bacteria may have therefore unintentionally developed resistance mechanisms against both in the past, which is why we may see sequences similar to synthetic compounds in ancient samples. For example, Aminocoumarins (Alt et al. 2011; Rahimi et al. 2016) and Spiropyrimidinetriones (Shi et al. 2019; Govender et al. 2022), although in slightly different ways, both target the DNA gyrase enzyme in bacteria. Another argument is that the cutoff was too relaxed, and the 50% sequence identity threshold resulted in FP Spiropyrimidinetriones and Sulfonamide ARGs. In the ancient and modern threshold analysis using simulated data, the FP rate for these specific ARGs increased as the threshold was loosened. Therefore, these synthetic ARGs may surface in ancient ARG analysis employing medium-low sequence identity thresholds. It is also possible that the decontamination protocol did not effectively remove these specific ARGs. However, this scenario is unlikely, as any sequences detected in environmental and laboratory controls would have been excluded from the calculus samples.

The large proportion of MLS ARGs is reasonable in ancient samples because these naturally manufactured antibiotics are produced by the 380 million-year-old microorganism *Streptomyces* (McDonald et al. 2017) a soil-borne bacteria isolated from a range of natural environments (Laskaris et al. 2010; Augustine et al. 2012; Encheva-Malinova et al. 2014; Zhao et al. 2016; Arocha-Garza et al. 2017; Sarmiento-Vizcaíno et al. 2018; Núñez-Montero et al. 2019; Nicault et al. 2020). If there are microorganisms that want to compete with *Streptomyces* in the natural world, *Streptomyces* may increase the antibiotic load in that environment, which would explain the occurrence of macrolide ARGs in ancient ecosystems. MSL ARGs have been previously observed in ancient dental calculus (Warinner et al. 2014; Rascovan et al. 2016) and their observation in other ancient samples may provide insight into how ancient lifestyles and diet influenced oral resistomes.

Since handwashing for hygiene was still in its infancy in the mid-1800s (Poczai and Karvalics 2022), post-industrial populations may have faced greater exposure to environmental bacteria, such as antibiotic-producing *Streptomyces*, when preparing food for example, potentially leading to a different form of selective pressure in the mouth compared to today. People in the 21^st^ century consume a Western diet containing pre-packaged, processed, high-sugar, high-fat, and fried meals (Clemente-Suárez et al. 2023) and it is widely accepted that sugary diets lead to oral disease (Echeverria et al. 2022; Guo et al. 2022; Echeverria et al. 2023; Spatafora et al. 2024), prompting the use of antibiotics that explain the high levels of resistance genes observed in our modern calculus. Additionally, antibiotics used to treat other diseases directly or indirectly linked to poor diets, along with antibiotics present in certain foods like meat, may also contribute to the resistance genes observed in modern calculus. It is clear that variations in lifestyle factors, such as diet and hygiene practices, have contributed to the prevalence of resistance to specific antibiotics in different periods (e.g., MLS in post-industrial samples and Tetracyclines today). This highlights the significant shifts in bacterial mechanisms within the oral microbiome over the past two centuries.

Moreover, learning how ancient diets impacted the oral resistome can help us gradually discover what types of foods and lifestyles may contribute to the emergence of AMR, which can assist in fostering healthier oral microbiomes today. For example, dental plaque is known to be a reservoir for AMR distribution (Anderson et al. 2023), so it can be argued that carbohydrate-rich eating habits and poor oral hygiene that help form plaque should be addressed to combat AMR. In this light, we should consider the oral microbiota a hotbed for ARGs.

Finally, we examine ancient MRGs in our post-industrial and modern dental calculus analyses. As humans became more industrialised and more metals contaminated the environment, including people’s noses, mouths, and lungs, we would expect more resistance to metals to emerge. Mercury and multi-metal ARGs were found in both ancient and modern samples. Before the Industrial Revolution, it would be normal for modest amounts of mercury to exert selective pressure on bacteria. Mercury is a naturally occurring contaminant that enters the environment through volcanic eruptions (Bagnato et al. 2009) and forest ecosystems (Zhou et al. 2021). Furthermore, metals have always been a part of human life, from cave fires to several metal ages, therefore the development of multi-metal resistant ARGs is not unexpected. Copper and Iron MRGs, absent in post-industrial dental calculus, make an appearance in modern calculus. This could indicate that modern industrialisation has contributed to the spread of AMR. Because ARGs and MRGs are co-selective agents, addressing AMR should consider the impact of pollution. To effectively quantify the influence of pollution on the spread of AMR, more data is needed to completely determine which MRGs rise over pre-industrial, industrial, post-industrial, and modern dental calculus.

## Conclusion

Our experiment using simulated ancient metagenomic datasets suggests that DNA damage has little effect on the quantity and type of ARGs recovered from ancient samples. In contrast, environmental contamination of ancient metagenomic datasets has a pronounced effect, artificially inflating the number and type of ARGs recovered. It is also clear that higher thresholds generate more TPs and TNs than FPs and FNs in comparison to medium-low thresholds. Therefore, ARGs are highly reliable at a ≥75% identity percentage threshold for ancient dental calculus when decontamination protocols are followed. High identity percentage thresholds can cause SNP errors in low-quality aDNA. As a result, medium to low thresholds may have to be used but certain drug-resistant groups are interpreted with caution. In conclusion, the principal findings of this study recommend appropriate thresholds for resistance gene analysis, emphasise the importance of effective environmental and laboratory decontamination processes and highlight the significance of using ancient and modern oral microbiomes as sources for ARGs and MRG analysis.

## Author contributions and funding information

Conceptualization: F.J.S, C.J.M. Supervision: A.T., C.J.M., and C.S. Funding acquisition: C.S., A.T., C.J.M. Investigation: F.J.S, G D-A., and C.J.M. Formal Analysis: F.J.S, G. D-A., C.J.M and A.T. Data Curation, Visualization and Writing-Original Draft: F.J.S. Writing - Review & Editing: all authors.

## Acknowledgements

The authors acknowledge the use of the University of Bradford High-Performance Computing Service in the completion of this work.

